# Mitotic chromosomes fold by condensin-dependent helical winding of chromatin loop arrays

**DOI:** 10.1101/174649

**Authors:** Johan H. Gibcus, Kumiko Samejima, Anton Goloborodko, Itaru Samejima, Natalia Naumova, Masato Kanemaki, Linfeng Xie, James R. Paulson, William C. Earnshaw, Leonid A. Mirny, Job Dekker

## Abstract

During mitosis, chromosomes fold into compacted rod shaped structures. We combined imaging and Hi-C of synchronous DT40 cell cultures with polymer simulations to determine how interphase chromosomes are converted into compressed arrays of loops characteristic of mitotic chromosomes. We found that the interphase organization is disassembled within minutes of prophase entry and by late prophase chromosomes are already folded as arrays of consecutive loops. During prometaphase, this array reorganizes to form a helical arrangement of nested loops. Polymer simulations reveal that Hi-C data are inconsistent with solenoidal coiling of the entire chromatid, but instead suggest a centrally located helically twisted axis from which consecutive loops emanate as in a spiral staircase. Chromosomes subsequently shorten through progressive helical winding, with the numbers of loops per turn increasing so that the size of a helical turn grows from around 3 Mb (~40 loops) to ~12 Mb (~150 loops) in fully condensed metaphase chromosomes. Condensin is essential to disassemble the interphase chromatin conformation. Analysis of mutants revealed differing roles for condensin I and II during these processes. Either condensin can mediate formation of loop arrays. However, condensin II was required for helical winding during prometaphase, whereas condensin I modulated the size and arrangement of loops inside the helical turns. These observations identify a mitotic chromosome morphogenesis pathway in which folding of linear loop arrays produces long thin chromosomes during prophase that then shorten by progressive growth of loops and helical winding during prometaphase.

**One Sentence Summary:** Mitotic chromosome morphogenesis occurs through condensin-mediated disassembly of the interphase conformation and formation of extended prophase loop arrays that then shorten by loop growth and condensin-dependent helical winding.

## Introduction

Chromosomes dramatically change their conformation as cells progress through the cell cycle. Throughout most of interphase, chromosomes are organized in a hierarchy in topologically associating domains (TADs) (1, 2) and A- and B-compartments (3). At a finer scale, chromatin looping between promoters, enhancers and CTCF-bound sites (4, 5) facilitates gene regulation. During mitosis, these features all disappear and chromosomes are compacted as compressed arrays of randomly positioned consecutive chromatin loops (6–9).

Although the organization of these two states is now increasingly understood, much less is known about how cells convert from one state into the other. Previous microscopy observations revealed that chromosomes become individualized during prophase and form linearly organized structures where sister chromatids are initially mixed (10–13). By late prophase, sister chromatid arms have split but remain aligned along their length. At this stage, each sister chromatid is thought to be organized as an array of loops that emanate from an axial core composed of condensin complexes, topoisomerase II alpha and possibly other proteins (14–18). This core may be composed of an association of individual complexes, rather than a single integrated structure (19). During subsequent prometaphase, the chromatids shorten and become thicker (11), ultimately producing the classical fully condensed metaphase chromosomes (20). How the compaction of loop arrays occurs during prometaphase is not known.

Here we employ a chemical-genetic system for highly synchronous entry of DT40 cells into prophase. This allowed us to apply Hi-C with high temporal resolution and to determine how chromosome conformation changes as cells disassemble the interphase nucleus, progress through prophase and prometaphase and form mitotic chromosomes (21, 22). These data, combined with polymer simulations (23, 24) and direct imaging of chromosomes reveal a mitotic chromosome morphogenesis pathway with distinct transitions in chromosome conformation, including compartment and TAD disassembly, loop array formation by late prophase and chromosome shortening during prometaphase through growing and winding of loops around a central helical scaffold. Using an auxin-inducible degron approach (25) we then identify distinct key roles for condensin I and II in this pathway.

## Results

### Synchronous progression into mitosis

To obtain cultures of cells that synchronously enter mitosis we arrested cells in G_2_ by selectively inhibiting CDK1, the kinase required for mitotic entry and progression. We stably expressed a variant of the *Xenopus laevis* CDK1 cDNA (CDK1as) harboring a F80G mutation in DT40 cells (21, 26). This mutation in the ATP binding pocket renders CDK1as sensitive to inhibition by the bulky ATP analog 1NM-PP1 (21). We then disrupted the endogenous CDK1 gene using CRISPR/Cas9. Growing cells for 10 hours in the presence of 1NM-PP1 efficiently arrested >90% of cells in G_2_ as indicated by FACS (**Table S1, Fig. S1**) and by microscopy analysis of chromosome and nuclear morphology (Fig. 1A). Washing out 1NM-PP1 from synchronized cells re-enabled ATP binding and led to rapid release of cells from the G_2_ arrest and synchronous entry into prophase.

**Fig. 1.**
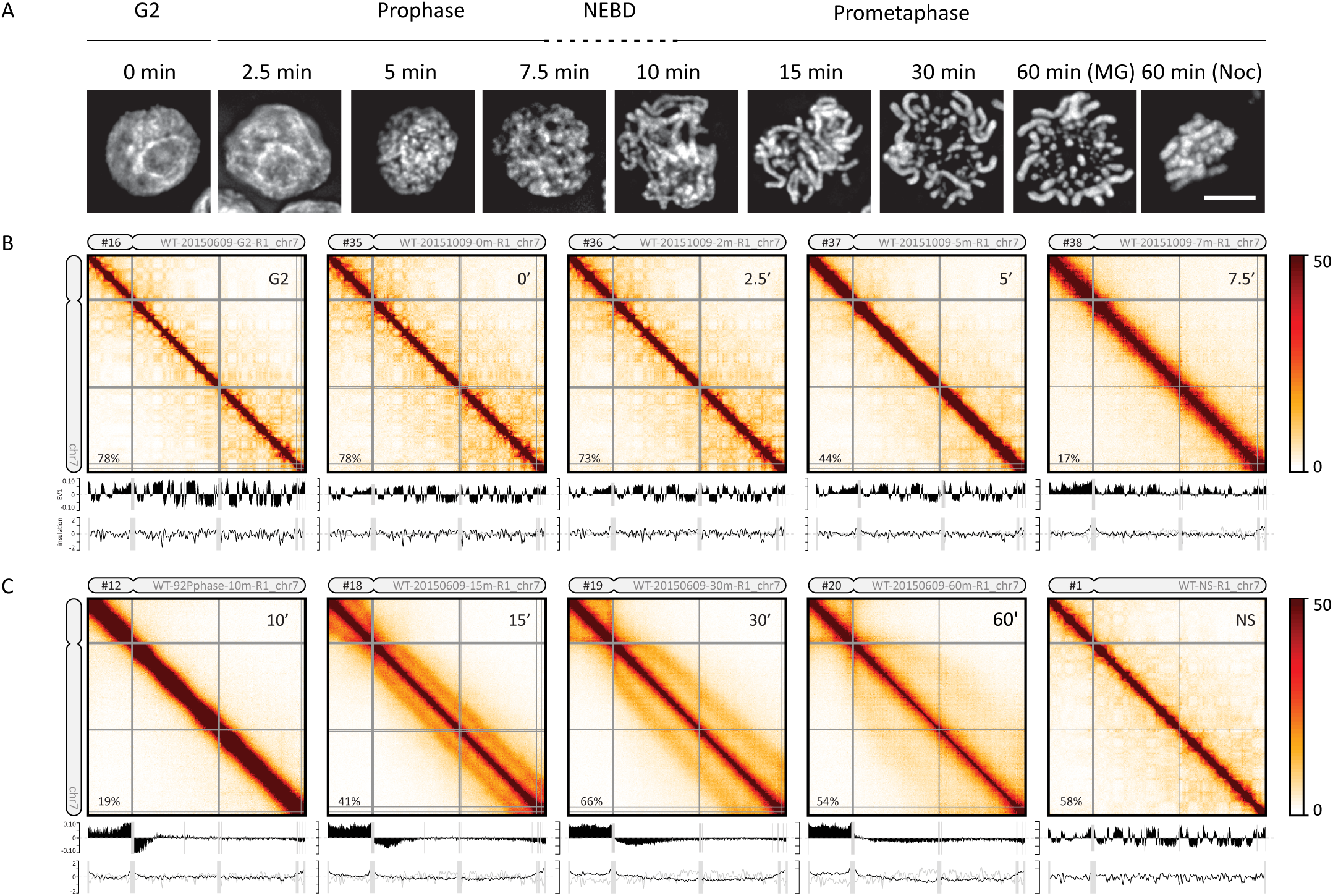
Chromosome morphogenesis during synchronous mitosis. (**A**) Representative DAPI images of nuclei and chromosomes in CDK1as DT40 cells taken at indicated time points (in minutes) after release from 1NM-PP1-induced G_2_ arrest. Bar indicates 5 micron. (**B-C**) Hi-C interaction maps of chromosome 7 (binned at 100 kb) from prophase (**B**) and prometaphase cells (NS = non-synchronous cells) (**C**). The numbers in the top left corner of the chromosome correspond to the sample numbers listed in **Table S2**. The top plot below each Hi-C interaction map displays compartment signal (Eigenvector 1; with percentage signal explained by first eigenvector). The bottom graph shows insulation score (TADs).

This cell system allowed us to study chromosome morphogenesis by harvesting cells at sequential time points for imaging and Hi-C analysis as they synchronously progress through mitosis. For cultures collected at later time points (30 - 60 minutes), we added nocodazole 30 minutes prior to their release from the 1NM-PP1 arrest, to block exit from metaphase into anaphase (see Methods). All time courses described here were performed in duplicate and results were highly concordant (Supplemental Material, e.g. **Fig. S6**). DAPI staining showed the expected chromosome condensation and individualization in prophase (Fig. 1A). Nuclear envelope breakdown (NEBD) occurred by around t = 10 minutes as evidenced by the disappearance of the smooth perimeter of the DAPI mass, by staining for Lamin B1, which diffuses into the cytoplasm upon NEBD (**Fig. S2**) (27) and by measuring the association of previously cytoplasmic condensin I subunits with the chromosomes (**Fig. S3B**). Previous studies, and our proteomic analysis (**Fig. S3B**) show that by late prophase, cohesin has dissociated from the arms of sister chromatids, which then separate, but remain aligned (11, 12, 28, 29). Chromosome shortening subsequently occurred during prometaphase and at the late time points fully condensed chromosomes were observed (Fig. 1A).

### Loss of compartments and TADs in Prophase

Hi-C analysis showed that G_2_-arrested cells display all features characteristic of interphase cells (8). First, chromosomes form territories as indicated by relatively high levels of intra-chromosomal interactions (3). Second, chromosomes display the characteristic compartmentalization into active A- and inactive B-compartments as revealed by the plaid pattern of Hi-C interactions (3) (Fig. 1B). The locations of A- and B-compartments in G_2_ resembled those detected in exponentially growing cells, though the compartment signal strength was stronger and the pattern sharper in the synchronous cells as a result of uniformity in cell cycle stage (compare Fig. 1B **(G_2_), C (NS)**). Third, TADs were readily visible in the Hi-C interaction maps as squares of relatively high interaction frequencies along the diagonal (Fig. 1B). TAD boundaries were similar in position and strength to those in non-synchronous cells as determined over a 250 kb window with the previously described insulation score (30) (**Fig. S4**). Finally, when the contact probability *(P)* was plotted as function of genomic distance (*s*) we found the characteristic inverse relationship previously observed for non-synchronous cells (**Fig. S5**). Together, these analyses show that G_2_ chromosomes, which are composed of two closely aligned and likely catenated sister chromatids, are organized similarly to G_1_ chromosomes (8).

This interphase chromosomal organization was rapidly lost upon release of cells into prophase. As soon as 5 minutes after release we detected a marked reduction in the typical plaid pattern of long-range interactions pointing to loss of compartments (Fig. 1B). By 10 minutes (late prophase) no compartments were detected. At the same time, TADs were also lost and the defined blocks of enriched interaction frequencies along the diagonal disappeared during prophase (Fig. 1B, **Fig. S6**).

We used eigenvector decomposition to quantify the disappearance of compartments (3). The first eigenvector readily captured compartments at t = 0 and 2.5 minutes, but starting at t = 5 minutes the first eigenvector explained progressively less of the variance in the Hi-C interaction maps, indicating weakening of the compartment structure (**Fig. S7**). By t = 7 minutes the strength of the first eigenvector was reduced to 17% and by t = 10 minutes, compartments were no longer captured by the first eigenvector. Loss of compartments was also quantified by calculating the ratio of A-to-A or B-to-B interactions over A-to-B interactions over the time course. From t = 0 to t = 2.5 minutes and onward this fraction decreases steadily, indicating that preferential interactions within compartments are lost (**Fig. S8**).

TAD boundaries have a low insulation score (high insulation), whereas for loci inside TADs this score is high (no insulation). The variance of the insulation scores can be used as a quantitative measure of the presence of TADs (8). Starting at t = 2.5 minutes the variance of the insulation profiles progressively decreased, indicating loss of TADs (Fig. 1B, Fig. 1C**, Fig. S4B**). By t = 7 minutes the variance was reduced more than 2-fold and by t = 10 minutes no TADs were detected. This disappearance of TAD boundaries was confirmed by directly calculating the average level of insulation at G_2_ boundaries during mitosis: insulation is the strongest in G_2_ and by late prophase insulation values are near background levels (**Fig. S9**). From these analyses we conclude that compartments and TADs disappear rapidly during early prophase.

By late prophase, when sister arms have resolved (11, 31), and around the time of nuclear envelope breakdown (t ~ 10 minutes), the main feature of the contact maps is a general inverse relationship between contact frequency and genomic distance that appears similar for all loci. This can be quantified by *P(s)* plots, which describe the probability of two sequences being crosslinked as a function of the distance between them (Fig. 2A). As prophase progresses the shape of *P(s)* changes: in G_2_ cells contact frequency decay is shallow up to a distance of several hundred kb, reflecting compaction within TADs (32, 33), but then for larger distances the decay becomes steeper. During prophase the initial shallow decay as *P(s) ~ s^−0.5^* continues for longer range interactions, with a steeper drop at 2 Mb at t = 10 minutes, suggesting a higher degree of compaction. Interestingly, the shape of the *P(s)* curve, and the decay of *P(s)* ~ *s^−0.5^* at late prophase resembles that of late prometaphase we described before (8). As we demonstrate below, this decay and shape are consistent with formation of a linearly arranged, layered organization of the chromosome (8), where the size of each layer corresponds to the position of a steep drop in the *P(s)* curve.

**Fig. 2.**
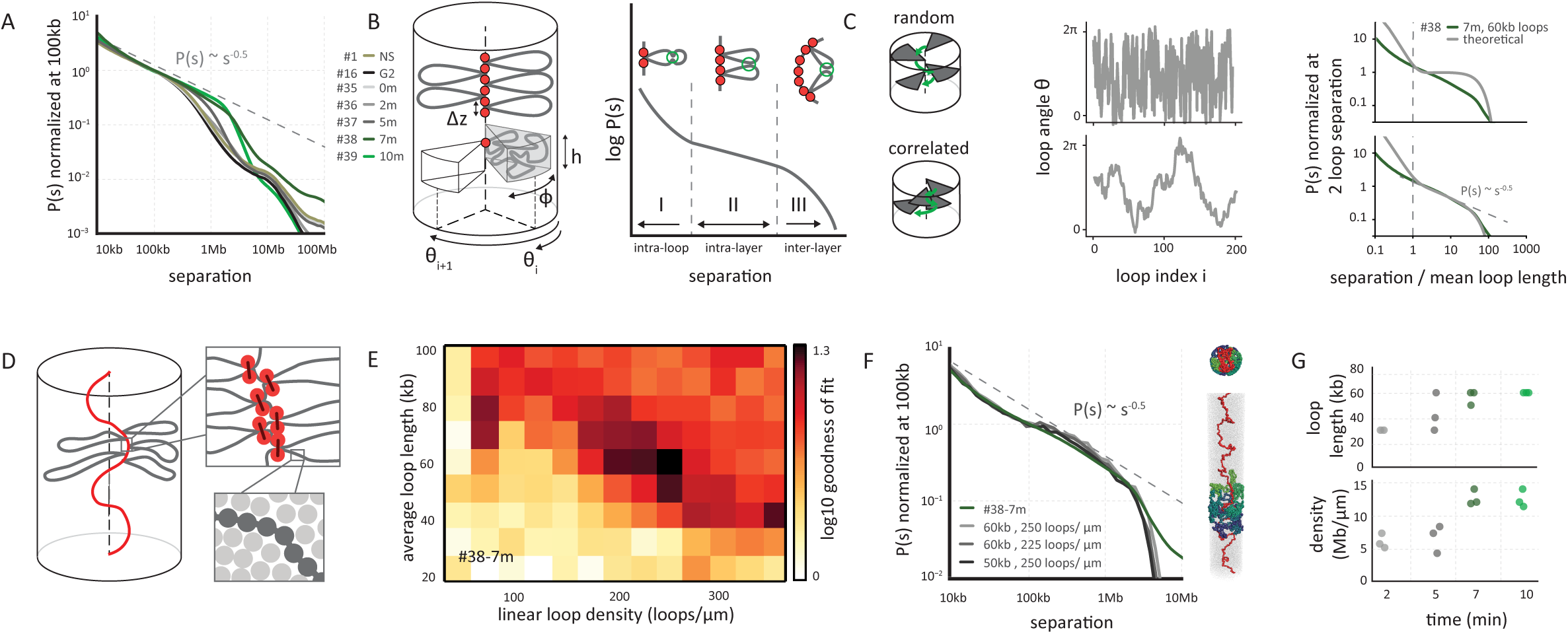
Prophase chromosomes fold as axially compressed loop arrays. (**A**) Genome-wide curves of contact frequency *P(s)* vs genomic distance *s*, normalized to unity at *s*=100 kb. The curves are derived from prophase Hi-C data at the indicated time points after release from G_2_ arrest. #numbers indicate the dataset identifiers listed in Table S2. The dotted line indicates *P(s)* = *s^−0.5^* observed for mitotic chromosomes *(8)*. (**B**) Overview of the coarse grained model of prophase chromosomes. The chromosome is compacted into a series of consecutive loops and compressed into a cylindrical shape. The loop bases form a scaffold at the chromosomal axis, each loop occupies a cylindrical sector of height h and angular size φ, oriented at angle Θ_i_. The coarse-grained model predicts the *P(s)* curve to have three distinct regions: an intra loop (I), intra layer (II) and inter layer (III) regions. (**C**) The best fitting *P(s*) predictions by the coarse grained model for late prophase (#38, t = 7 minutes) under two different assumptions on loop orientations: (top panels) random uncorrelated and (bottom panels) correlated orientations of consecutive loops. Uncorrelated angular loop orientations lead to a plateau in *P(s)* in the intra-layer, whereas correlated angles lead to the experimentally observed *P(s)* = *s^−0.5^* (right panels). (**D**) Overview of polymer simulations of prophase chromosomes. Chromatin fibers are modeled as chains of connected particles (dark grey circles), compacted into arrays of consecutive loops (loop bases indicated in orange). The chromosomes are compacted into a cylinder with a particle density of one nucleosome per 11x11x11nm cube (lower right). (**E**) Goodness of fit for simulated vs experimental *P(s)*. Polymer simulations were performed for a range of linear loop densities and average loop length, and for each simulation *P(s)* was calculated. The heatmap shows the quality of a match between the predicted and experimental *P(s)* curves at late prophase (#38, t = 7 minutes) for each parameter set. (**F**) *P(s)* derived from late prophase Hi-C experiments (green line) and the best fitting polymer models (grayscale lines). The legend shows the average loop size and linear density of loops along the chromosome axis in the corresponding models. On the right an example of a model is shown with loops bases in red and several individual loops rendered in different colors. (**G**) The average loop size and linear density of the 3 best-fitting models of prophase chromosomes at different time points.

### Appearance of a second diagonal band in Hi-C maps from prometaphase cells

At t = 15 minutes, when cells have entered prometaphase, the Hi-C maps display a *P(s)* curve with a drop at 2 Mb. Strikingly, a distinct second diagonal band that runs in parallel with the primary band is observed for all loci and chromosomes (Fig. 1C). This second diagonal represents increased interaction frequencies between any pair of loci separated by around 3 Mb. At 15 min, this feature is clearly observed in *P(s)* plots as a local peak at ~3 Mb (Fig. 3A**, Fig. S10A**). As cells progress through prometaphase the position of the drop in *P(s)* and the position of the second diagonal move to larger genomic distances (Fig. 1C; Fig. 3A; **Fig. S10A**). By t = 60 minutes, when compact metaphase chromosomes have formed, the second diagonal is positioned at ~12 Mb and appears more diffuse.

**Fig. 3.**
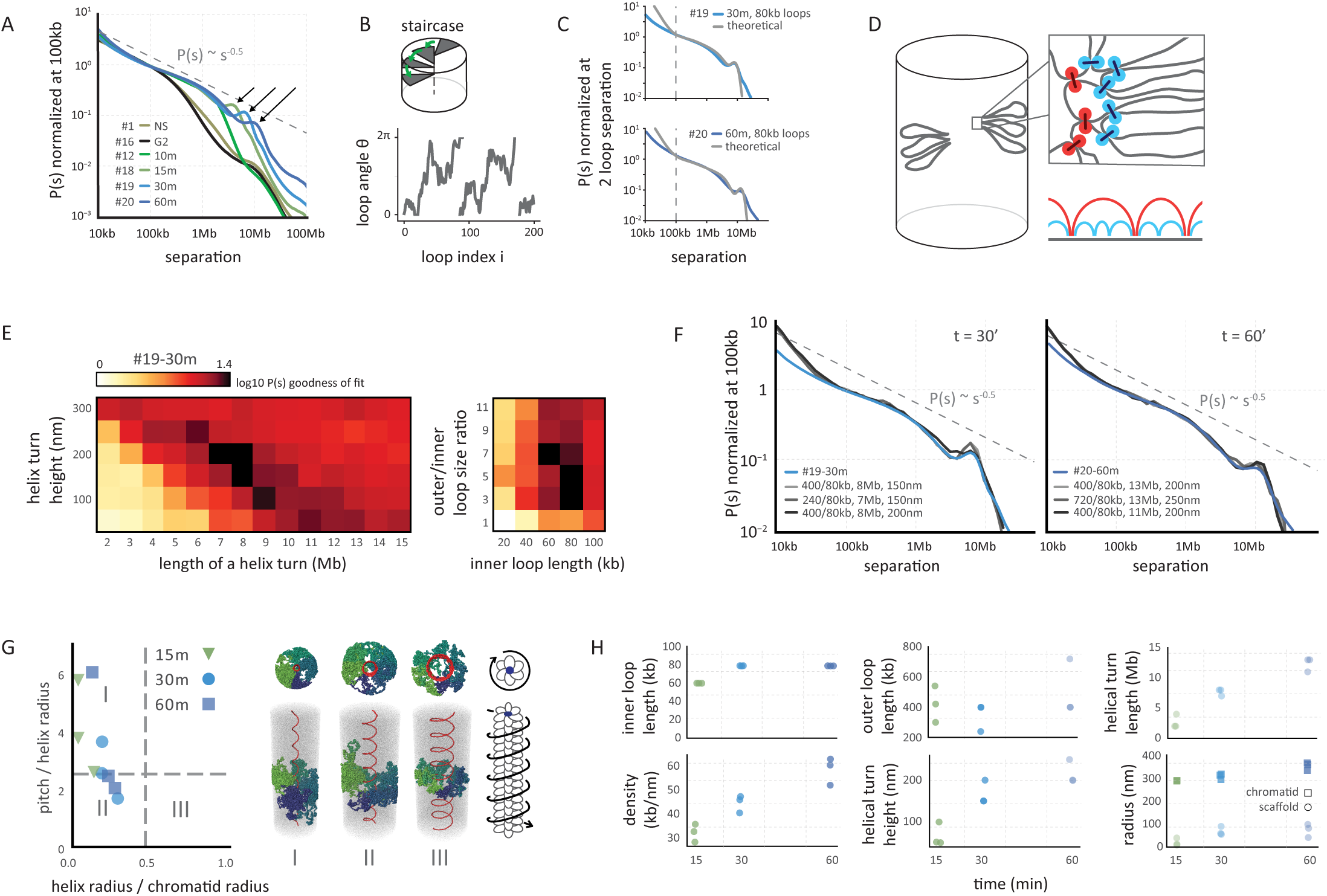
Helical organization of prometaphase chromosomes. **(A)** Genome-wide curves of contact frequency *P(s)* vs genomic distance (separation, *s*), normalized to unity at *s*=100 kb. The curves are derived from Hi-C data obtained from prometaphase cells (t = 10-60 minutes after release from G_2_ arrest). The dashed line indicates *P(s)* = *s^−0.5^*. Arrows indicate positions of a local peak in *P(s)* representing the second diagonal band in Hi-C interaction maps. **(B)** The coarse grained model of prometaphase chromosomes with staircase loop arrangement. Top: the staircase loop arrangement implies that loops rotate in genomic order around a central scaffold (see Supplemental Materials). Bottom: angles of adjacent loops are correlated and steadily increasing, reflecting helical arrangement of loops. **(C)** The best fitting *P(s)* predictions by the staircase coarse grained model for late prometaphase t = 30 minutes (#19-30m; top panel) and t = 60 minutes (#20-60m, bottom panel) after release from G_2_ arrest (Hi-C data: blue lines; model; gray lines). **(D)** Overview of polymer simulations of prometaphase chromosomes. Chromosomes are modeled as arrays of consecutive nested loops with a helical scaffold (outer loops in red, inner loops in blue, also indicated diagrammatically bottom right). **(E)** Goodness of fit for simulated vs experimental *P(s)*. Polymer simulations were performed varying the helix height (nm), the size of a helical turn (Mb), and the sizes of inner and outer loops. For each simulation *P(s)* was calculated. The heatmaps show the quality of the best match between the predicted and experimental *P(s)* at prometaphase (#19, t = 30 minutes), when two out of four parameters were fixed to the specified values. **(F)** *P(s)* derived from prometaphase Hi-C experiments (blue lines) and the best fitting polymer models (gray lines). Left panel: t = 30 minutes (#19-30m), right panel t = 60 minutes (#20-60m) after release from G_2_ arrest. The legend shows the average size of outer and inner loops, the length of a helix turn and the helical pitch in the corresponding models. **(G)** The geometric parameters of the helical scaffolds from the best fitting polymer models of prometaphase chromosomes. The x-axis shows the ratio of the radius of the helical scaffold to that of the whole chromatid and indicates how broad the helix appears; the y-axis shows the ratio of the pitch to the helix radius and indicates how axially extended the chromosome appears. The dashed lines show the corresponding values (0.46 and 2.5122) for the optimal space-filling helix (*80*). Classical solenoid configurations are predicted to be in sector III, while “spiraling staircase” configurations are in I and II. On the right three examples of models of type I, II and III are shown with loops bases in red and several individual loops rendered in different colors. Also shown is a schematic of a prometaphase chromosome with the helical winding of loops indicated by arrow around the loop array. **(H)** The parameters of the best 3 models of prometaphase chromosomes at different time points.

Such a location-independent non-monotonic behavior of the *P(s)* curves indicates a periodicity of interactions that reflect chromosome structure at the scale of megabases. Under defined experimental conditions, chromosomes also exhibit structures called bands along the length of their arms (34). However, bands are irregular in size. Another explanation could be the formation of multi-Mb sized loops with bases along a central chromosome axis, but this would lead to an unrealistic morphology of extremely short, dense chromosomes; for instance, a 20 Mb chromosome would constitute only a handful of such loops leading to a very short axis the length of which would only be several condensin complexes.

The only known regular periodic structural feature of chromosomes is helical coiling, which has been observed in certain fixed chromosome preparations (10, 35–37). When chromosomes are helically organized with a helical pitch (the length of a complete turn) of ~3 Mb, each locus is in relatively close proximity to loci located one turn up or down the chromosome, i.e. 3 Mb up or downstream along the DNA. The progressive movement of the second diagonal band to larger distances during prometaphase would then reflect an increased size (in Mb) of the helical turn.

Although this is the first Hi-C data revealing the helical coiling of chromosomes, the chicken DT40 late prometaphase (t = 60 minutes) Hi-C interaction maps strongly resemble those for mitotic human HeLa S3 chromosomes that we had reported earlier (8). In addition, we have re-analyzed mitotic HeLa S3 Hi-C data in more detail by deeper sequencing. This also revealed a weak second diagonal band at ~10 Mb distance (**Fig. S11**), suggesting this periodic folding is a conserved feature of mitotic chromosomes.

### Testing models of chromosomes

Previous studies suggested that prophase and mitotic chromosomes are organized as arrays of consecutive loops emanating from a condensin-rich scaffold, forming a polymer bottlebrush (38, 39), with layered organization of loops (6, 20, 40). To classify and interpret *P(s)* curves of Hi-C data during prophase and prometaphase we therefore started by building coarse-grained models of chromosomes as arrays of loops. In our coarse-grained models, a chromosome is represented by a cylinder with the scaffold located at the cylindrical axis; loop bases are arranged consecutively along the scaffold, and each chromatin loop, emanating from the scaffold in a particular direction, is represented by a blob of loci that belong to this loop (Fig. 2B, (see Supplemental Material). Loops are regularly placed along the axis of the chromosome, their angular positions are determined by a specific stochastic model; loop sizes are exponentially distributed and bases of loops are not positioned at defined genomic sequences or loci (8). For specific models of loop arrangements, presented below, the *P(s)* curve can be found analytically as the return probability of a stochastic process describing angular positions of loops (see Supplemental Material, section “Coarse-grained model of contact probability decay in mitotic chromosomes”). The resulting *P(s)* always has three regions (Fig. 2B): (i) an intra-loop region at short separations, where two loci are likely to be within the same loop, reflecting the internal organization of loops; (ii) an “intra-layer” region at larger genomic separations that reflects the specific arrangement of loops relative to each other within a radial layer of the cylindrical chromosome, and (iii) an “inter-layer” region following a relatively steep drop in contact frequency at large genomic distances, where loci are located in loops that are so distant along the scaffold that their blobs can no longer overlap. In the *P(s)* plot of experimental Hi-C data throughout mitosis the intra-layer region and the drop-off can be readily discerned (Fig. 2A, 2C, 2F).

### Prophase chromosomes

The coarse-grained models show that the relative orientation of consecutive loops strongly affects the shape of the *P(s)* curve in the intra-layer region. If loops emanate from the axis in random directions, i.e. the orientations of consecutive loops are independent of each other, the contact frequency *P(s)* does not decay with genomic distance in the intra-layer region, as any pair of loops within a layer are equally likely to interact (Fig. 2C). In contrast, introducing correlations between orientations of consecutive loops, i.e. forcing neighboring loops to project in similar directions, makes them follow an angular random walk, which produces a distinct power-law decay of *P(s)*~s^−0.5^. The angular random walk is a 1D random walk on a circle and has a return probability of *P(s)*~s^−0.5^ until the full turn is made by the walk. The *P(s)*~s^−0.5^ decay followed by a drop is in good agreement with the late prophase Hi-C data (t = 7-10 minutes – Fig. 2C). Taken together these results suggest that by late prophase chromosomes are already organized into arrays of consecutive loops with correlated angular orientations.

To gain quantitative insights into the size, arrangement, and density of loops for late prophase chromosomes, we developed detailed polymer models. In these models chromatin is represented as a 10 nm fiber (41, 42), where one monomer corresponds to one nucleosome (Fig. 2D), allowing us to simulate up to 40 Mb of chromatin. Prophase chromosomes are modeled as arrays of consecutive loops of exponentially distributed length and random genomic locations, emanating from a flexible scaffold, as would result from a loop extrusion process (43). The array of loops is further condensed by imposing poor solvent conditions to the density observed in electron microscopy (one nucleosome per 11x11x11nm cube, i.e. ~40% volume fraction) (44), while preserving the overall cylindrical shape of the chromosome (Fig. 2D). We systematically varied two parameters of the model: the average loop size and the linear loop density along the chromosomal scaffold (Fig. 2E). For all combinations, we generated equilibrium conformations, simulated a Hi-C experiment, and evaluated its ability to reproduce *P(s)* curves from Hi-C data for different time points during prophase (Fig. 2E-G).

We found that polymer models can accurately reproduce *P(s)* (20 kb<s<4 Mb) for all prophase time points, in agreement with the prediction of the coarse-grained architecture with correlated orientations of consecutive loops (Fig. 2C). The best matching models for later prophase time points, when sister chromatids are separate and run side-by-side, have gradually increasing average loop size: from 40-50 kb at t = 5 minutes to ~60-70 kb at t = 10 minutes (Fig. 2G), reproducing the gradually shifting position of the drop-off from 2 to 3.5Mb (i.e. increase of the layer size), while maintaining about the same ~50 loops per layer and ~250 loops per μm. These results are consistent with a model where loop arrays are formed at early time points in mitosis, and loop sizes grow gradually, e.g. by merging smaller adjacent loops (24). We conclude that both coarse grained modes and polymer simulations indicate that by late prophase, chromosomes fold as dense arrays of loops, with consecutive loops positioned with correlated radial orientations.

### Prometaphase spirals

A striking feature of prometaphase Hi-C data is the appearance of the second diagonal band, which manifests itself as a distinct peak on the *P(s)* curves (Fig. 3A, arrows). This feature indicates a periodicity of chromosome structure at the scale of megabases. A possible explanation for the second diagonal band could be the presence of interactions between sister chromatids. Throughout mitosis, sister chromatids are accurately aligned (45) but the degree to which sister chromatids are mixed changes from high in early prophase to low in prometaphase when sisters are essentially resolved (11, 12). The second diagonal band, on the contrary is absent in prophase and becomes more prominent through prometaphase, suggesting that it cannot emerge due to interactions between sisters. To directly test whether interactions between sister chromatids can give rise to the second diagonal we simulated pairs of aligned prophase chromosomes, representing pairs of sisters, that overlap to different extents and found that regardless of the extent of overlap, a second diagonal band is never observed (Supplemental Methods, **Fig. S12A**). We conclude that the second diagonal observed in Hi-C data is unlikely to result from interactions between aligned sister chromatids.

As argued above, periodic interactions seen by Hi-C are most readily explained by a helical organization of mitotic loop arrays, which has been observed microscopically (6, 10, 37, 46). Two classes of chromosome architecture can give rise to periodicity in contact frequencies: an “external” helix when the whole chromosome is folded into a solenoid (46) (the solenoid model), and a “staircase” model in which consecutive loops wind in a helical order around a centrally located scaffold (“internal” helix). We note that by “scaffold” we do not necessarily imply a solid integrated structure stretching from one end of the chromosome to the other. The “scaffold” could equally be a dynamic association of smaller complexes that pack with helical symmetry. By modeling we examined these classes of architectures and the continuum of models in between them.

To explore whether an internal helix can arise in the dense array of loops through reorganization of loop orientations, while preserving the cylindrical morphology of the whole chromosome, we first extended our coarse-grained prophase model (Fig. 2B) by adding a preferred angular orientation for each loop: (1) as in prophase, the orientation of each loop is correlated with its neighbors; (2) in addition, loops have preferred, but not fixed, orientations that follow a helical path, thus winding around the chromosomal scaffold (Fig. 3B). Loops in this spiral staircase model follow an angular Ornstein-Uhlenbeck random walk with bias toward preferred positions, and *P(s)* can be found analytically (47) (Supplemental Material, section “Coarse-grained model of contact probability decay in mitotic chromosomes – loops with spiral staircase orientation”). This coarse-grained model yields a *P(s)* curve that closely follows the experimental prometaphase *P(s)* and displays both the *P(s)~s*^−0.5^ decay and the narrow peak corresponding to the second diagonal band (Fig. 3C). These results indicate that, (i) the emergent second diagonal band in Hi-C data can result from a spiral organization, and (ii) such organization can arise from preferred orientations of loops around the central scaffold.

Detailed polymer modeling allowed us to explore a broader range of architectures, with both external and internal helices, and to obtain quantitative estimates of loop sizes and other aspects of organization. Two aspects of the prometaphase organization must be captured by any model: (i) a higher linear density of chromatin of up to 50-70 Mb/μm necessitating an evolution of the loop architecture; and (ii) spiraling of the scaffold. The higher density of loops can be achieved by a nested loop organization where several smaller (inner) loops are organized consecutively within each larger (outer) loop whose bases form the central axis (Fig. 3D). We found that the presence of nested loops is an essential feature for prometaphase models, as models with a single layer of loops cannot reproduce Hi-C *P(s)* curves even when other parameters were varied (**Fig. S12B**). To model helical architecture we made the scaffold follow a helical path in 3D, while allowing loops to adopt their equilibrium conformations within an otherwise cylindrical chromosome (Fig. 3D).

We systematically varied the parameters of the model, such as geometry of the spiral scaffold and loop sizes (Fig. 3E,F), which also probed different lengths and widths of chromosomes as the volume density was kept constant. We found that for t = 30 minutes the best agreement was achieved for the scaffold forming a relatively narrow internal spiral (R=30-60nm) approaching the staircase architecture (Fig. 3G). This spiral is much more narrow than the ~300nm diameter of the chromatid, and has a small pitch (the height of one turn, 100-200nm) (Fig. 3H). Interestingly, such a spiral arrangement of loop bases is sufficient to achieve helical winding of loops that reproduces the second diagonal in the interaction maps and the peak on the *P(s)* curves for t = 15, 30 and 60 minutes (Fig. 3F-H). Wider spiraling of the scaffold (Fig. 3GIII) approaching external helix architectures (46) failed to accurately reproduce *P(s)* (**Fig. S12C**). Taken together, coarse-grained and polymer models that agree with Hi-C data overwhelmingly support the spiral staircase architecture of the scaffold (I and II in Fig. 3H), which in turn provides helical winding of loops.

Fitting consecutive time points probed by Hi-C, we found that the linear chromatin density of the best-matching models continued to grow throughout prometaphase, which agreed with the observed steadily shortening of mitotic chromosomes (Fig. 1A, 3H, **Fig. S13**). This increase in linear density reflects the shift of the drop-off in the *P(s)* curves to larger genomic distances for t = 15 to 30 to 60 minutes. Simulations show that shifting the peak in *P(s)* to larger genomic distances representing the second diagonal in consecutive time points (Fig. 3A) can be achieved by increasing the radius of the helical scaffold from 30 to 100 nm, the radius of the chromatid from 300 to 360 nm, and increasing the pitch from 100 to 250 nm, while maintaining a constant outer and inner loop size (~400 kb and ~80 kb respectively) (Fig. 3H). These changes lead to the increase in amount of DNA (Mb) per turn of the spiral from ~3 Mb in early prometaphase up to ~12 Mb by late prometaphase.

### Condensins are critical for prophase chromosome morphogenesis

To determine the role of condensin complexes in chromosome morphogenesis, we depleted both condensin I and II together in Cdk1as DT40 cells by fusing an auxin inducible degron domain (AID) to the SMC2 core subunit (Supplemental Materials, **Table S3**). In the presence of the plant F-box protein osTIR1 (driven by a CMV promoter in these cells) addition of auxin induces rapid proteasome-dependent degradation of the SMC2-AID protein, thus disrupting both condensin I and II complexes (18, 25, 48). Cells were arrested in G_2_ by a 10 hour incubation in the presence of 1NM-PP1 followed by an additional 3 hours in the presence of auxin and 1NMPP1 (Supplementary Materials) after which SMC2 levels were reduced to <5% (**Fig. S14**). Depletion of SMC2 in G_2_-arrested cells did not affect global chromosome organization as compartments and TADs were comparable to those in WT G_2_ arrested cells (Fig. 4A, **Fig. S15**). Cells entered prophase rapidly after washout of 1NM-PP1, and the onset of NEBD, as indicated by DAPI staining, occurred as in wild type at ~10 minutes (**Fig. S16**).

**Fig. 4.**
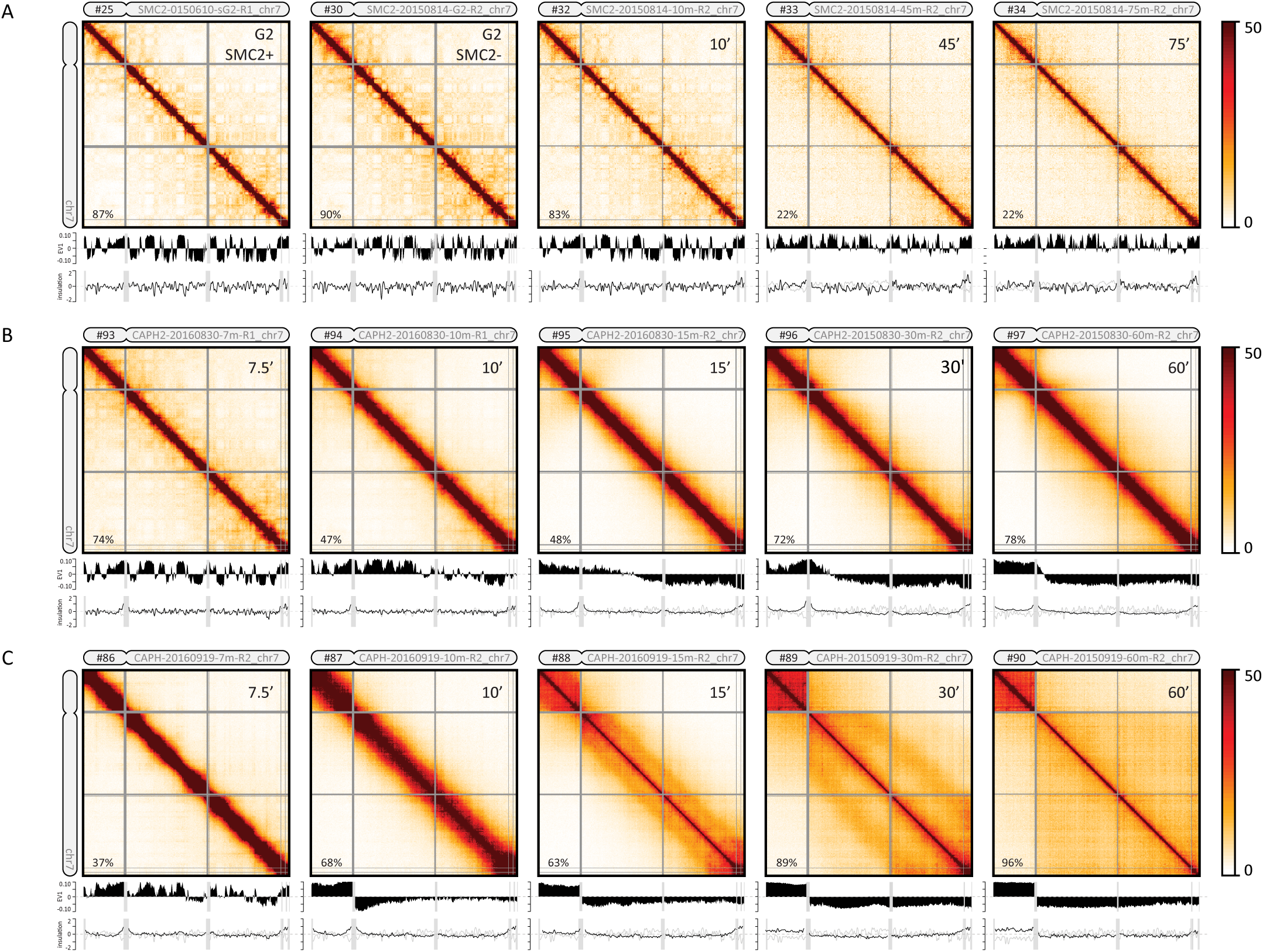
Defects in chromosome morphogenesis in condensin mutants. (**A-C**). Heatmaps showing Hi-C interaction frequency maps (binned at 100 kb) for chromosome 7 at indicated time points (top right in each heatmap) after release from G_2_ arrest. The first plot below each Hi-C interaction map displays the compartment signal (Eigenvector 1). The bottom graph shows the insulation score (TADs; binned at 50 kb). The numbers in the top left corner of the chromosome correspond to the data set identifiers in **Table S2**. (**A**) SMC2-AID cells were treated with auxin for three hours prior to release from G_2_ arrest to deplete SMC2. SMC2+: Hi-C interaction map for G_2_-arrested cells prior to auxin treatment. SMC2-: Hi-C interaction map for G_2_-arrested cells after three hours of auxin treatment. (**B**) Hi-C data for CAP-H2-AID cells treated for three hours with auxin prior to release from G_2_ arrest to deplete CAP-H2. (**C**. Hi-C data for CAP-H-AID cells treated for three hours with auxin prior to release from G_2_ arrest to deplete CAP-H.

Microscopy analysis showed that chromosomes in SMC2-depleted cells did not form well-resolved chromatids as cells progressed to prometaphase, confirming previous observations (**Fig. S16**) (18, 49–51). Chromatin in such cells is lacking functional condensin (52, 53), but nonetheless manages to achieve a normal degree of chromatin compaction despite the absence of individualized chromosomes (54) despite the absence of individualized chromosomes (*54*). FACS analysis confirmed that these cells are incapable of exiting M-phase and entering G_1_ subsequently, even after 3 hours from G_2_ release (**Fig. S1**).

Hi-C analysis revealed that in the absence of SMC2, interphase compartments and TADs were still present and largely unaffected by late prophase, at a time when they were completely disassembled in WT (t = 10 minutes, Fig. 4A, **Fig. S15A**). Furthermore, individualized prophase chromosomes were not observed by microscopy (**Fig. S16, Table S1**). NEBD did occur; indicating cells progressed to physiological prometaphase. In prometaphase (t = 45 minutes and t = 75 minutes), compartments and TADs became progressively weaker, but remained detectable (Fig. 4A, **Fig. S17-19**). No second diagonal, characteristic for WT prometaphase (Fig. 1C) ever appeared in Hi-C contact maps (Fig. 4A), instead *P(s)* curves show little change from G_2_ (**Figs. S15A, S20-22**). The reduction in compartment strength was quantified by plotting average interaction frequencies between loci arranged by their eigenvector values. When we quantified A-to-A interactions and B-to-B interactions separately we found progressive loss of compartment signal, but a weak A-to-A signal remained detectable even at late time points (**Fig. S15A, S18A**). Analysis of the variation of the insulation score along chromosomes indicated that TAD boundaries were reduced in strength but not eliminated (Fig. 4A, **Fig. S19**). Further, removal of cohesin (SMC1/3) and CTCF from chromatin, as assessed by chromatin enrichment for proteomics (ChEP) (*55*) was delayed and reduced compared to WT (**Fig. S3**). This may explain the incomplete loss of TAD boundaries. Combined, these data reveal that condensin is not required for TAD and compartment architecture during interphase. In its absence, mitotic chromatin is condensed but chromosomes do not become individualized or acquire the normal mitotic morphology, while partially preserving elements of interphase architecture. This indicates (i) that condensation and formation the rod shape of mitotic chromosomes are two separate processes, supporting assumptions of our model; and (ii) the critical role for condensin complexes is in the formation of proper morphology and internal organization of mitotic chromosomes, and in disassembly of the interphase architecture (*56*).

### Condensin I and II play distinct roles in chromosome morphogenesis

Next, we determined the roles of condensin I and II separately. We fused auxin inducible degron domains to the condensin II-specific kleisin CAP-H2 (CAP-H2-AID) or the condensin I-specific kleisin CAP-H (CAP-H-AID) in CDK1as DT40 cells (Supplemental Methods). Cells were arrested in G_2_ by incubation with 1NM-PP1 (**Fig. S1**) and CAP-H-AID or CAP-H2-AID degradation was induced by addition of auxin leading to >95% protein depletion (**Fig. S14**). Cells were then released from the G_2_ block and chromosome conformation was determined by microscopy and Hi-C as cells progressed through mitosis. Depleting either condensin I or II led to less severe phenotypes than depleting both together (SMC2-AID): degradation of either CAPH-AID or CAP-H2-AID did not prevent cells from progressing through mitosis with typical changes in chromosome conformation, including the appearance of individualized chromosomes during prometaphase (**Fig. S16**). In contrast to cells lacking both condensin I and II (SMC2-AID), these cells were able to exit mitosis within 3 hours after entry into prophase (**Fig. S1**).

Comparison of Hi-C interaction matrices (Fig. 4BC, **Fig. S15BC**) and *P(s)* curves (Fig. 5A, 5B, **Fig. S20BC, S21-22**) for CAP-H and CAP-H2 depleted cells in late prometaphase (t = 30 minutes and 60 minutes) shows that they closely reproduce distinct parts of the WT *P(s)*, capturing different aspects of the WT architecture. The *P(s)* curve for CAP-H2-depleted cells, where only condensin I remains active, matches that of the intra-layer organization of WT up to ~6 Mb, and did not indicate the presence of a second diagonal band (Fig. 5B). The *P(s)* curve for CAP-H depleted cells (active condensin II), in turn matches that of WT only for the long-range organization (6-20 Mb), including the second diagonal band (Fig. 5B). CAP-H depleted cells have a much lower contact frequency between loci separated less than 6 Mb as compared to WT and CAP-H2 deleted cells. These observations suggest condensin I and II play distinct roles, and at different structural levels, in mitotic chromosome morphogenesis, providing a mechanistic explanation for earlier microscopic studies (*57–60*).

**Fig. 5.**
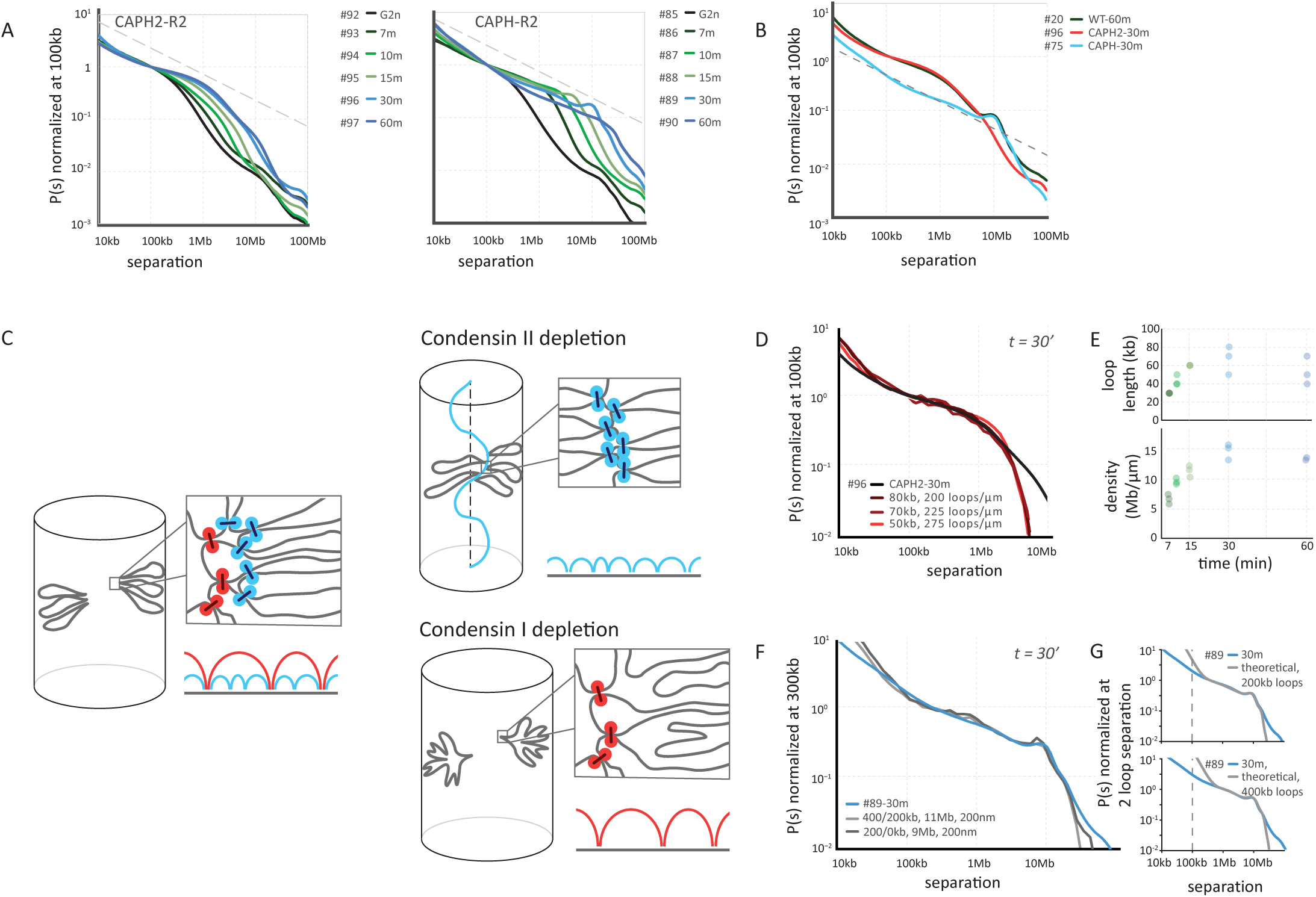
Distinct roles for condensin I and II in helical folding of prometaphase chromosomes. **(A)** Genome-wide curves of contact frequency *P(s)* vs genomic distance *s*, normalized to unity at *s*=100 kb. The curves are *P(s)* derived from Hi-C data obtained from CAP-H2-depleted (left panel) and CAP-H-depleted cells (right panel), at t = 7-60 minutes after release from G_2_ arrest. Dashed line indicates *P(s)* = *s^−0.5^*. (**B**) The effect of CAP-H and CAP-H2 depletion on the experimental *P(s)* curves in prometaphase. (**C**) Overview of polymer simulations of CAP-H2 (top) and CAP-H (bottom) depleted chromosomes. Top: in our polymer model of prometaphase chromosomes, removal of CAP-H2 is modeled via removal of outer loops and relaxation of the helix, effectively reverting the chromosomes to prophase architecture. Bottom: removal of CAPH is modeled via removal of the inner loops, while preserving the helical arrangement of the scaffold. Condensin II loop anchors are shown in red, condensin I loop anchores are shown in blue. (**D**) *P(s)* derived from late prometaphase CAP-H2 depletion Hi-C experiments (black line) and the three best fitting polymer models (red lines). The legend shows the average loop size and linear density of loops along the chromosome axis in the corresponding models. (**E**) The average loop size and linear density of the 3 best-fitting models of CAP-H2-depleted chromosomes at different time points. The legend shows the size of the loops (kb) and the number of loops per μm (linear density). (**F**) *P(s)* derived from late prometaphase CAP-H depletion Hi-C experiments (blue line) and the best fitting polymer models with and without nested inner loops (grayscale lines). The legend shows the average size of outer and inner loops, the length of a helix turn and the helical pitch in the corresponding models. (**G**) The best fitting *P(s)* predictions by the staircase coarse grained model for late prometaphase CAP-H depletion Hi-C experiments at t = 30 minutes (#89-30m) after release of G_2_ arrest (blue lines). Top: assuming the loop size of 200 kb, bottom: the loop size of 400 kb.

### Helical winding during prometaphase requires condensin II

In condensin II-depleted cells, chromosomes lose their interphase conformation as cells enter prophase. Both A- and B-compartments and TADs were lost starting around the prophase-prometaphase transition (t = 10 - 15 minutes; Fig. 4B, **Fig. S15B; Fig. S17, Fig. S18**). In late prometaphase (t = 30 - 60 minutes), chromosomes in CAP-H2 depleted cells were longer and narrower than WT chromosomes, as has been observed before >*5*>,>*6*>,>*60*</i>)(<i>59</i>, <i>61</i>, <i>60</i>)(<i>59</i>, <i>61</i>, <i>60</i>) (**Fig. S16**). *P(s)* curves for t = 10 and 15 minutes (early prometaphase) resembled those in WT for late prophase (t = 10 minutes; compare Fig. 5A with Fig. 2A), displaying a mild decay followed by a steep drop that is characteristic for a densely packed loop array (Fig. 2B). Most strikingly, depletion of CAP-H2 prevented emergence in prometaphase of the second diagonal band in Hi-C contact frequency maps and *P(s)* plots (Fig. 4B, 5B; **Fig. S15B, S20**).

The close similarity between CAP-H2 prometaphase and WT prophase Hi-C, and the lack of the second diagonal, prompted us to model CAP-H2 chromosomes as a prophase-like array of a single layer of loops emanating from a flexible, non-helical scaffold. By systematically varying the loop size and the degree of linear compaction, we found that excellent agreement with experimental *P(s)* curves was achieved for ~40-60 kb loops, and a linear density of 15 Mb/μm for all prometaphase time points (Fig. 5D, 5E). This linear density is 3-4x smaller than that of WT prometaphase chromosomes (50-70 Mb/μm). These simulations reproduce the long and narrow chromosomes observed by microscopy, indicating that in the absence of condensin II, chromosomes form extended prophase-like loop arrays and do not progress to further longitudinal shortening and helical winding during prometaphase.

### Condensin I modulates the internal organization of prometaphase helical layers

Cells depleted for CAP-H (**Fig. S14**) seemed to progress through prophase normally: Hi-C data show a rapid loss of compartments and TADs (Fig. 4C, **S15C, S18-19**) and by late prophase individual chromosomes could be discerned by DAPI staining (**Fig. S16**). Deviation from the WT morphogenesis pathway was observed during prometaphase, i.e. after NEBD, when the bulk of condensin I normally loads in WT (**Fig. S3B, C**). A second diagonal was observed at 30 minutes indicating helical winding of the chromatids (Fig. 4C, **S15C**) but this diagonal was located at a genomic distance of ~12 Mb, which in WT cells was only observed at t = 60 minutes. This indicates that the progression to larger helical turns as prometaphase progresses is accelerated in cells lacking CAP-H.

Despite similar spiral organization, loss of condensin I leads to a different arrangement of loops, and different folding of individual loops, as seen from differences in the *P(s)* curves: the intra-layer arrangement of loops shows a characteristic *P(s)~s^−0.5^* from 400 kb to ~3 Mb, with *P(s)* for the *s*<400 kb region having a different slope, possibly reflecting a different intra-loop organization. We found that this intra-layer regime, the second diagonal band, and the drop-off *P(s)* curve are captured well by the coarse-grained model with 200-400 kb loops emanating with correlated angular orientations from the spiral scaffold (Fig. 5G). This loop size agrees well with the sizes of outer loops in the best models for WT chromosomes at t = 60 minutes (Fig. 3H), suggesting that in the absence of CAP-H (condensin I) chromosomes do not form smaller inner loops, while maintaining larger outer loops (Fig. 5C).

Polymer simulations allowed us to test whether removal of inner loops in the WT model can reproduce Hi-C data for CAP-H depleted cells. Strikingly, when we matched the t = 30 minutes *P(s)* curve with the simulations of prometaphase chromosomes with helical scaffolds and nested loops, the best match was achieved with either a single layer of 200 kb loops, or a nested system of loops, with 400 kb outer loops and 200 kb inner loops (Fig. 5F). These results show that loss of condensin I results in the loss of the 60-80 kb inner loops while maintaining ~200-400 kb outer loops emanating from a helical staircase scaffold. Together these results suggest that CAP-H (condensin I) is essential for formation of short (60-80 kb) loops but is dispensable for formation of the helical arrangement of the scaffold.

Taken together, our data obtained with CAP-H and CAP-H2 depleted cells support the model of formation of nested loops during prometaphase. If the larger, outer loops are formed by condensin II and smaller, inner loops by condensin I, then depletion of CAP-H2 (condensin II) should result in the loss of outer loops with a remaining single layer of condensin I-mediated smaller inner loops. Loss of CAP-H (condensin I) should result in the loss of inner loops with a remaining layer of condensin II-mediated larger outer loops. Both predictions are consistent with our polymer simulations (Fig. 5).

## Discussion

### A mitotic chromosome morphogenesis pathway

The data and modeling presented here suggest a chromosome morphogenesis pathway by which cells convert interphase chromosome organization into compacted mitotic chromosomes (Fig. 6). Together, our imaging and Hi-C data, coarse-grained models and polymer simulations, and previous observations (11) reveal that upon entry into prophase, interphase features such as compartments and TADs are disassembled within minutes in a condensin-dependent process and by late prophase, chromosomes are organized as radial loop arrays.

Our models that achieve best agreement with Hi-C data show that during prophase, condensin II-dependent loops grow from 30-40 kb to 60 kb in size, leading to a ~2-fold increase in linear chromatin density from ~7 Mb/μm to 15 Mb/μm. Condensins at loop bases form a chromosomal scaffold (*20, 61*), which may be a dynamic, rather than static structure, and loops are arranged consecutively along it (one loop every ~5 nm of the axis). Interestingly, the radial arrangement of loops around the central flexible scaffold is not random, with consecutive loops projecting in similar directions i.e. with an angularly correlated arrangement.

**Fig. 6.**
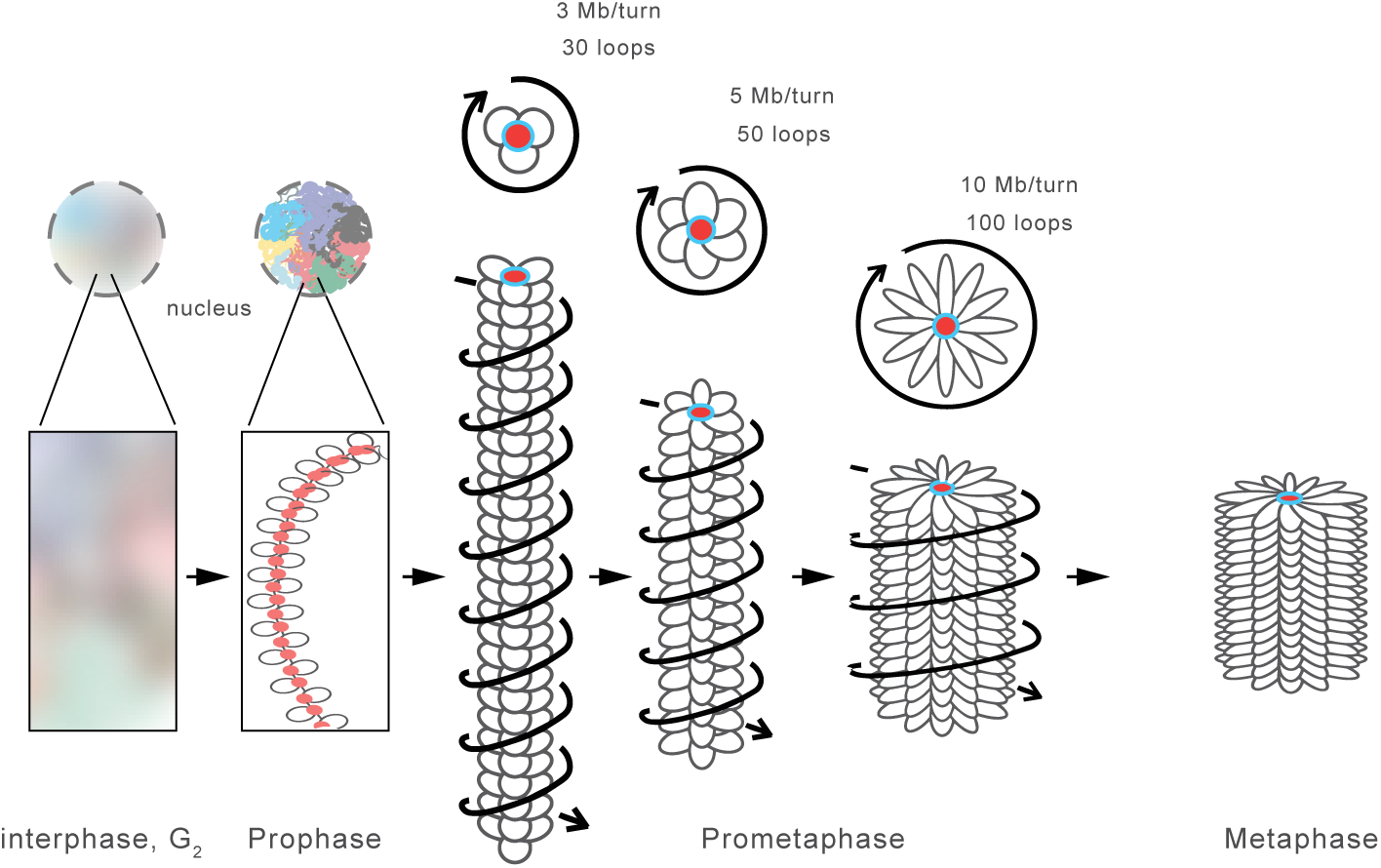
A mitotic chromosome morphogenesis pathway. Model for prophase and prometaphase conformation. As cells enter prophase interphase features such as compartments and TADs are disassembled. In prophase, condensin II generates long loop arrays and sister arms separate. The scaffold of condensin II-mediated loop bases is indicated in red. Upon nuclear envelope breakdown and entry into prometaphase condensin II generates larger loops that are split in smaller ~80 kb loops by condensin I. Chromosomes are shown as arrays of loops (only inner loops can be observed microscopically; top: cross-section, bottom: side view). The nested arrangements of centrally located condensin II-mediated loop bases and more peripherally located condensin I-mediated loop bases are indicated in red and blue respectively. At this time the central scaffold acquires a helical arrangement with loops rotating around the axis as in a “spiral staircase” (helical path of loops is indicated by arrows). As prometaphase progresses outer loops grow and the number of loops per turn increases and chromosomes shorten to form the mature mitotic chromosome.

Chromosomes shorten along their longitudinal axis and become wider during prometaphase. Our simulations show that condensin II loops continue to grow to 200-400 kb by 30 min and 400-700 kb by 60 min, accompanied by an increase in the linear chromatin density, which reaches 60 Mb/μm. However, two important reorganizations take place during prometaphase. First, large condensin II-mediated loops are subdivided into smaller 80 kb loops in a condensin I-dependent process, thus producing a nested loop arrangement with ~400 kb outer loops and ~80 kb inner loops. Second, the loop array acquires a helical arrangement as evidenced by the appearance of a second diagonal band in Hi-C maps for all loci and chromosomes. Models show that this helical arrangement of loops can be achieved if the scaffold forms a narrow helical “spiral staircase” inside an otherwise homogeneous cylindrical chromosome. Interestingly, the period, radius and pitch of this helix continue to grow through prometaphase. An emerging model of the prometaphase chromosome thus has a central helical scaffold formed by condensin II (*61*) and supporting 200-400 kb outer loops, that are further subdivided into 80 kb condensin I-mediated inner loops, emanating from the scaffold in correlated orientations and further condensed into a high volume density.

### Comparison to previous and classical studies

While specific details of this model emerge from an unbiased fitting of models to the data, the emerging organization and its quantitative characteristics agree with earlier studies. First, the 60-80 kb sizes of the inner loops, which unlike the outer loops can be directly observed microscopically, are remarkably similar to (1) earlier experimental measurements of 80 kb loops based on electron microscopy (6, 20); (2) to an estimate of 60 kb loops based on an extensive survey of the literature (40) and (3) to 80-100 kb loops inferred from Hi-C analysis of mitotic HeLa cells (8). Similarly, changes of linear density from prophase to prometaphase in the best models (from 15 Mb/μm to 50 Mb/μm) are consistent with prophase chromosomes being at least two-fold longer than metaphase chromosomes (*11, 58*).

Second, helical prometaphase chromosomes have been observed in certain fixed chromosome preparations (10, 35, 37), and this has led to a number of different models for how mitotic chromosomes are folded. One set of models proposes that mitotic chromosomes are hierarchically folded as coiled coils to ultimately form a ~250 nm thick fiber that then coils to form the mature mitotic chromosome (*62, 63*). Other models include the formation of radial loop arrays and propose these arrays can coil to form helically arranged chromosomes (37, 46). Our analysis of Hi-C data indicates that the prometaphase chromosome is organized around a helical central region or scaffold: loops emanate with helical packing from a centrally located “spiral staircase” scaffold. Modeling shows that other helical arrangements of loop arrays, e.g. coiling of the entire loop array itself as proposed by Rattner and Lin (46) are not consistent with our Hi-C data.

Our helical scaffold/loop model unifies a range of models and observations made over the years. First, the model includes loop arrays, for which there is extensive evidence (6, 8, 20, 40), as well as helical features of chromosomes, observed by microscopy. Our model also explains how a helical chromatin packing arrangement can be achieved while scaffold proteins such as condensins and topoisomerase II are localized centrally, as has been documented before (15–17), within a cylindrical chromatid that is not obviously helical when visualized with a DNA dye such as DAPI. Consistent with our models, Maeshima and Laemmli (*64*) observed a centrally located distribution of condensins and topoisomerase II twisted like a barber pole, and those authors also proposed that loop arrays and helical folding both occur together. Interestingly, by late prometaphase we estimate the height of one helical turn to be around 200 nm, which is also the size of the layer (12 Mb layer at linear density 60 Mb/um), and is consistent with microscopic observations by Strukov et al. (45) suggesting that consecutive genomic loci follow a helical gyre with a pitch of ~250 nm within the cylindrical shape of chromatids.

### Possible mechanisms

We argue that such loop arrangements can naturally emerge due to the loop extrusion process. Loop extrusion as a mechanism of chromosome compaction has been hypothesized (*65*) and most recently examined by simulations (*24, 43, 66*) and supported by single-molecule studies (*67*). In this process, each condensin starts forming a progressively larger loop until it dissociates or stops because the DNA movement is blocked by neighboring condensins or other DNA-binding proteins. A recent study demonstrated that this process can form an array of consecutive loops (8) with condensins forming a central scaffold in the middle of a cylindrical chromosome (43), essential features of mitotic chromosomes. Formation of extruded loops along each chromatid in cis, in combination with topoisomerase activity, can also drive sister chromatid resolution (43). We note that sister chromatids are resolved by late prophase (11–13) indicating that the formation of loop arrays occurs as sister chromatid arms become separated.

Another aspect of loop extrusion is that loop sizes are established by a dynamic process of condensin exchange, without a need for barrier elements or specific loading sites (24). This is consistent with our Hi-C data that suggests that loop bases are not positioned at specific reproducible positions (e.g. scaffold or matrix attachment regions - (*68, 69*)) in a population of cells. If loops were positioned at reproducible positions this would have led to the appearance of sharp boundaries and locus-specific pairwise interactions between loop anchors in Hi-C interaction matrices (8). We found Hi-C interactions to be similar for all loci, indicating a lack of positional preference for prophase loops, and in agreement with formation of loops by condensin-mediated extrusion. Furthermore, simulations show that loop extrusion slowly approaches steady state by exchanging condensins and gradually increasing loop sizes during this process (24). This is consistent with gradual growth of loops up to 500 kb by slowly-exchanging condensin II, and relatively rapid formation of 60-80 kb inner loops by the more rapidly exchanging condensin I (*70*).

The rate of loop growth by condensin II appear to be about the same in prophase 7Kb/min (70Kb in 10min) and prometaphase ~8Kb/min (~500Kb in 60min), and remarkably consistent with recent single-molecule measurements (bioRxiv https://doi.org/10.1101/137711). If loop extrusion is performed by two connected motors progressing in opposite directions, the speed of loop extrusion is twice, the speed of each motor (measured as 3.6Kb/min), resulting in 7.6Kb/min.

Formation of nested loops was critical for our polymer simulations to reproduce Hi-C data because it allowed a higher linear chromatin density. In this architecture, only outer loop bases are located at the central scaffold, while the loop bases of inner loops are radially displaced. The presence of two types of loops is further supported by the distinct roles of condensin I and II: in the absence of condensin I outer loops remain, while in the absence of condensin II, arrays of smaller loops are formed, indicating that condensin II generates outer loops, and condensin I generates inner loops. Preliminary simulations show that arrays of nested loops can be formed by two types of loop extruders differing in residence time. Furthermore, we note that in prophase, when only condensin II is associated with chromosomes, there is a single array of loops, while in prometaphase, when condensin I gains access to chromatin, nested loops form. Thus, loop extrusion can provide a mechanism by which arrays of loops with specific average loop size but random positions form, leading to the formation of a condensin II scaffold. Subsequently, during prometaphase, condensin I gains access to chromatin and starts to generate nested loops.

Correlated orientations of loops and formation of a helical scaffold, however, requires some additional mechanisms. One possibility is that condensins located at the bases of loops interact to organize the orientation of loops (*61, 71*). Such interactions between condensins can in principle also lead to formation of the helical scaffold, which could be either static or dynamic. Why condensin II-based scaffolds only acquire helicity in prometaphase, and not in prophase is not known, but this could involve interactions with other proteins, such as DNA topoisomerase IIalpha or KIF4A. Our estimates of the radius of the prometaphase scaffold of 30-100 nm is consistent with a 50 nm length of SMC coiled coils that can interact with each other through HEAT repeats (*72*) which are known for ability to self-assemble into a helical “spiral staircase” (*73*). Gradual formation of such a HEAT-mediated staircase and binding of other factors can explain how the pitch and the radius of the helix increase in time.

We note that mitotic chromatin still condenses in the absence of both condensin I and II, although individualized rod-shaped chromosomes are not formed and cells cannot progress into anaphase. This indicates that there are other mechanisms by which chromatin fibers become condensed during mitosis. Our simulations also show that to achieve agreement with Hi-C data, chromatin should also be condensed (computationally analogous to poor solvent conditions) forming densely packed chromatin loops within mitotic chromosomes analogous to the dense packing of chromatin observed in mitotic chromosomes by electron microscopy (*41, 74, 75*). The molecular basis for this condensation is not known but may involve mitosis-specific chromatin modifications (*76, 77*) or active motor proteins such as KIF4A (*78, 79*).

The chromosome morphogenesis pathway described here, and the identification of distinct architectural roles for condensin I and II in organizing chromosomes as nested loop arrays winding around a helical “spiral staircase” within a cylindrical chromatid can guide future experiments to uncover the molecular mechanisms by which these complexes, and others such as topoisomerase IIalpha and KIF4A, act in generating, (re-)arranging and condensing chromatin loops to build the mitotic chromosome.

## Acknowledgements

Supported by grants from the National Human Genome Research Institute (HG003143) and the National Institutes of Health Common Fund (DK107980) to J. Dekker and L. Mirny. J. Dekker is an investigator of the Howard Hughes Medical Institute. Work in the Mirny lab was also supported by the National Science Foundation (Physics of Living Systems, 1504942), and the National Institute of General Medical Sciences (GM114190). Work in the Earnshaw lab was funded by Wellcome, of which W.C.E. is a Principal Research Fellow (Grant 107022). The Wellcome Centre for Cell Biology is supported by core funding from the Wellcome Trust [203149]. M.T.K. was supported by JSPS KAKENHI Grant (16K5095), and research grants from Mochida Memorial Foundation for Medical and Pharmaceutical Research, SGH Foundation, The Sumitomo Foundation, and The Canon Foundation. We thank Dr. Hakan Ozadam for assistance with Hi-C data processing. All Hi-C data has been submitted to GEO and will be publicly available upon publication.

